# A non-coding role for trypanosome *VSG* transcripts in allelic exclusion

**DOI:** 10.1101/2025.03.21.644657

**Authors:** Douglas O. Escrivani, Sebastian Hutchinson, Michele Tinti, Jane E. Wright, Catarina A. Marques, Joana R. C. Faria, Anna Trenaman, David Horn

**Affiliations:** Wellcome Centre for Anti-Infectives Research, School of Life Sciences, University of Dundee, Dow Street, Dundee DD1 5EH, UK; Quantum-Si France, 38 Rue De Berri 75008 Paris, France; School of Life Sciences, University of Dundee, Dow Street, Dundee DD1 5EH, UK; Centre for Parasitology, School of Infection and Immunity, University of Glasgow, UK; Biology Department and York Biomedical Research Institute, University of York, UK

**Keywords:** Antigenic variation, cncRNA, monoallelic, *Trypanosoma brucei*

## Abstract

Bloodstream-form African trypanosomes display mono-telomeric expression of a Variant Surface Glycoprotein (VSG) gene in an inter-chromosomally bridged transcription and splicing compartment, such that the dominant gene produces 10,000 times more transcript than excluded *VSG* genes. Antigenic variation, whereby parasites switch to express other VSGs, then underpins a robust host immune evasion strategy. Specific chromatin and RNA-associated factors are required to maintain *VSG* exclusion, but our understanding of the mechanisms involved remains incomplete. Here we show that the *VSG* transcript impacts allelic competition. We induced either specific translation blockade by recruiting MS2 coat protein to the active *VSG* 5’-untranslated region, or *VSG* transcript depletion using RNA interference. Neither perturbation substantially compromised exclusion of native *VSGs*, as determined by transcriptomic analyses. In contrast, exclusion of a *VSG* transgene was compromised when the native transcript was transiently depleted. Notably, while both perturbations blocked cytokinesis, an additional round of DNA replication and mitosis was observed when the transcript, known to be stabilized by a bloodstream-form specific cyclin-like F-box protein, was translationally blocked. We conclude that the *VSG* transcript is a bi-functional coding and non-coding RNA that participates in allelic competition to establish exclusion.

**Significance statement:** Allelic exclusion mechanisms underpin immune evasion in parasites and olfaction in mammals but the mechanisms responsible remain mysterious. VSG exclusion factors have been identified in trypanosomes, while RNA has been implicated in olfactory receptor exclusion, and in *var* gene exclusion in the parasites that cause malaria. The current study demonstrates a role for RNA in VSG exclusion in trypanosomes.

## INTRODUCTION

Trypanosomatids are flagellated protozoa and include several vector-transmitted parasites that impact both human and veterinary health. The African trypanosome, *Trypanosoma brucei*, is transmitted by tsetse flies, and causes lethal human and animal diseases. *T. brucei* is exclusively extracellular and presents a paradigm for studies on antigenic variation, which underpins evasion of host adaptive immune responses (1,2). In preparation for transmission to a mammalian host, *T. brucei* activates Variant Surface Glycoprotein (VSG) expression in the tsetse fly salivary gland, where cells initially express multiple *VSGs* prior to establishing monoallelic expression (3). The active and many silent bloodstream-form *VSG* expression sites are subsequently stably inherited, with estimates of switching frequency consistently occurring in substantially less than 1 % of cells per generation (4,5).

The VSG is a super-abundant and essential protein, and each bloodstream-form cell surface is coated with approximately ten million copies. Indeed, the active *VSG* gene accounts for approximately 10 % of the total cellular mRNA and protein. Typically, a single active *VSG* is transcribed by RNA polymerase I in an extranucleolar compartment known as the Expression Site Body or ESB (6), although two simultaneously active *VSGs* can also share an ESB (7). Despite the presence of fifteen competent, promoter-associated, polycistronic, and telomeric *VSG* expression sites in the 427-strain used here (8), the active *VSG* produces ∼10,000 times more mRNA than silent *VSGs* (9); transcription is initiated at all expression site promoters but is attenuated at excluded sites (10). This extreme form of transcriptional dominance and monogenic expression operates in the context of an (inter-chromosomal) RNA polymerase I transcription and splicing compartment that integrates a telomeric *VSG* and an RNA *trans*-splicing locus (11–13).

Several proteins contribute to maintaining monogenic *VSG* expression. These include positive regulators of *VSG* transcription, ESB1 (14) and SUMO (15), and CFB2, a cyclin-like F-box protein which binds and stabilizes *VSG* transcripts (16,17). The VSG exclusion (VEX) complex (18,19) is required to maintain exclusion, and forms an inter-chromosomal protein bridge that connects the (VEX2-associated) *VSG* transcription and (VEX1-associated) splicing sub-compartments (12,13). Also required to maintain exclusion are the telomere and RNA-binding protein, RAP1 (20,21), and its interaction with PIP5Pase (22). In addition, the histone tri-methyltransferase, DOT1B is required to rapidly silence inactivated *VSGs* (23,24) and both the chromatin chaperone, CAF-1 (18,25), and cohesin (26) promote stable inheritance of the active *VSG;* CAF-1 does so by binding the VEX-complex (18).

Super-abundant *VSG* mRNA incorporates a highly conserved ‘16-mer’ sequence in its 3’-untranslated region (UTR), and this motif has been implicated in binding RAP1 and antagonizing RAP1-based silencing (20), in binding CFB2 (17), and in promoting *N*^6^-methyladenosine (m^6^A) modification in the poly-A tail (27); both m^6^A and CFB2 stabilize the mRNA. Transcription of a second *VSG* driven by T7 phage RNA polymerase induces silencing of the active *VSG* (23), but *VSG* mRNA knockdown fails to induce activation of silent *VSGs* (28). Specific transcription blockade at the active *VSG* locus does induce activation of silent *VSGs* (29), however, consistent with a maintenance mechanism involving transcription-dependent sequestration of the limiting VEX-complex (12), or ESB1 (14), at the *VSG* transcription and splicing compartment.

Despite substantial advances in our understanding of monogenic *VSG* expression control in recent years, our understanding of how the known regulators detailed above establish and maintain *VSG* transcriptional dominance remains incomplete. To further explore allelic competition, and specifically to address the role of the *VSG* transcript, we generated strains in which translation of the active *VSG* could be conditionally and transiently blocked. Using native and transgene *VSG* expression assays, we compared these strains to *VSG* mRNA knockdown strains. Using the transgene assay, we identified a specific exclusion defect associated with transient transcript knockdown, indicating that establishment of *VSG* exclusion is *VSG* transcript dependent.

## MATERIALS AND METHODS

### Trypanosome cell culture

Wild-type, 2T1 (30) and derivative bloodstream-form Lister 427 cells were cultured in HMI-11 medium supplemented with 10 % fetal bovine serum (not heat inactivated) at 37 °C with 5 % CO_2_. 2T1 cells were genetically manipulated using cytomix, and a nucleofector II (Lonza) with 0.2 mm cuvettes (Bio-Rad), as previously described (31). MCP^VSG-2^, and RNAi*^VSG-2^*cells were subcloned and checked for the expected severe growth-defect prior to analysis. For the *VSG-5* transgene assay, parental 2T1, MCP^VSG-2^, and RNAi*^VSG-2^* cells were induced with tetracycline for 3 hours. Cells were then transfected with VSG-5 reporter (19). Cells were washed three times with HMI-11 to remove any residual tetracycline and selected with geneticin (G418) for 5 days. Antibiotic selection was applied at 10 µg.ml^−1^ blasticidin, 2 µg.ml^−1^ geneticin, 1 µg.ml^−1^ puromycin or phleomycin, and 5 µg.ml^−1^ hygromycin-B. Inducible expression systems were activated using tetracycline, which was applied at 1 µg.ml^−1^.

### Plasmid construction

The MCP tandem dimer (tdMCP) sequence was obtained from Addgene plasmid #40649 (phage-ubc-nls-ha-tdMCP-gfp) (32). The sequence was PCR-amplified using the primers, MS2CP-LaNLS-HA-HindIII-F (CCCC*AAGCTT*ATG<u>CGAGGACACAAGCGGTCACGTGAA</u>TACCCCTACGACGTGCCCGACTAC GCC) and MS2GFPR (GCTA*GGATCC*TTACTTGTACAGCTCGTCCATGC). The SV40-NLS sequence at the N terminus of MCP was replaced with a *T. brucei* La-NLS (underlined in primer and encoding RGHKRSRE (33); to generate pMCP^GFP^. This fragment was cloned in pRPa^xGFP^ (34) using the HindIII and BamHI sites (italics). A *BLA* selectable marker cassette was PCR-amplified with FWD_BLA_MS2 (CTAGT*GGATCC*TCTAGATGGGTCCCATTG) and with the *VSG-2* start codon within an *Sph*I site (italics) and an MS2 hairpin sequence (35) (underlined) and a portion of the β-tubulin 5’-UTR in the reverse primer, REV_MS2_BLA (GAAG*GCATGC*TGTTCTCCAGTTTTGTGTT<u>CTTAAGGCCTGATGGTCCTTAAG</u>TAGATAATT

TCGACTATTTTCTTTGATGAAAG). The PCR product was digested with BamHI and SphI (italics) and ligated to a sequence targeting the *VSG-2* gene, digested with BglII and SphI to generate pVSG^MS2^. The MCP^GFP^ and VSG^MS2^ constructs were digested with AscI and XhoI / HindIII, respectively, prior to sequential transfection, to first generate an MCP^GFP^ strain and then to generate the MCP*^VSG-2^* strains. The stem-loop VSG RNAi construct was derived from the pRPa^iSL^ vector (34). A ∼500 bp fragment of the *VSG-2* gene was PCR-amplified using the primers, *VSG-2SL-F* (GATCTCTAGAGGATCCGAGGAGCTAGACGACCAAC) and *VSG-2SL-R* (GATCGGGCCCGGTACCATAGTGACCGCTGCAGAAA).

### *VSG-2* RNA analysis

RNA was isolated from whole cell extracts using the Qiagen RNeasy kit. Reverse transcription of mRNA was performed using M-MLV reverse transcriptase (Promega). First strand cDNA synthesis of *VSG* was primed with a ‘3’-UTR’ primer, DH3 (GACTAGTGTTAAATATATCA), while PCR was with DH3 and a ‘spliced leader’ primer, SL22 (GAACAGTTTCTGTACTATATTG). Sanger sequencing was performed using either a spliced leader primer or a *VSG-2* specific primer. Northern blotting was performed using standard protocols. 2 µg of total RNA was run on a reducing (1.2 % formaldehyde) agarose (1.5 %) gel and stained with ethidium bromide. A ∼500 bp *VSG-2* fragment was radio-labelled with ^32^P dCTP (Perkin Elmer) using the large Klenow subunit and random primers for 15 min at 37 °C. Probes were hybridized overnight. We used a storage phosphor screen and visualized blots using a Fujifilm FLA-500 image reader.

### Protein blotting

For western blot analysis, 1 × 10^7^ cells were lysed and solubilized in 1× SDS sample buffer containing 0.1 M DTT at 55 °C for 20 min. Proteins were resolved by SDS-PAGE (approximately 5 × 10^5^ cell equivalents/lane) on NuPAGE bis-Tris 4 % to 12 % gradient acrylamide gels (Invitrogen) and transferred to nitrocellulose membrane (Invitrogen). Expression of MCP^GFP^ and EF1α was detected using a polyclonal sheep α-GFP antibody (MRC-PPU, 2-238, 1:10,000) and mouse α-EF1α (Millipore, CBP-KK1,1:10,000) in blocking buffer (50 mM Tris-HCl pH 7.4, 0.15 M NaCl, 0.25 % BSA, 0.05 % (w/v) Tween-20, 0.05 % NaN_3_ and 2 % (w/v) Fish Skin Gelatin). Detection was performed using IRDye 800CW donkey anti-goat (1:15,000) and IRDye 680RD donkey anti-mouse (1:10,000), respectively, in blocking buffer. The immunoblot was analyzed on the LI-COR Odyssey Infrared Imaging System (LICOR Biosciences).

### Metabolic labeling

For metabolic labeling, 10^7^ cells were collected by centrifugation (1,000 g, 10 min), washed and re-suspended in methionine and cytosine depleted RPMI and dialyzed fetal bovine serum (Thermo Fischer Scientific) for 15 min prior to labeling. Cells were then labeled with 50 µCi/ml ^35^S methionine (Perkin Elmer) for 5 min, washed in PBS and re-suspended in NuPAGE® LDS sample buffer (ThermoFisher) at 70 °C for 10 min. Samples were run on 8-12 % gradient polyacrylamide gels (Life Technologies). Gels were vacuum dried using a 583 gel-dryer (Bio-Rad). ^35^S incorporation was visualized on film after exposure at -80 °C.

### Microscopy

Cells were fixed in 1 % paraformaldehyde and attached to 12-well 5 mm slides (Thermo Scientific) by drying overnight. Following rehydration in PBS for 5 min, cells were blocked with 50 % FBS in PBS for 15 min. After two washes in PBS, cells were incubated in primary antibody for 1 h at RT; rat α-VSG-2 (1:10,000) and rabbit α-VSG-5 (1:10,000). Following three washes in PBS, cells were incubated in secondary antibody for 1 h at RT; α-rat Alexa 488 (Life Technologies, 1:2000), α-rabbit Alexa 568 (Life Technologies,1:2000). Cells were washed three more times in PBS and mounted in Vectashield with DAPI (4’,6-diamidino-2-phenylindole). Cells were imaged as z-stacks (0.1–0.2 µm) at 63 x magnification with oil immersion and a Zeiss Axiovert 200 M microscope with Zen Pro software (Zeiss). Images were processed using Fiji v1.5.2e (36). MCP^GFP^ fluorescence was directly visualised. For cell cycle analysis, cells were rehydrated in PBS and stained with DAPI (4’,6-diamidino-2-phenylindole) in Vectashield (Vector Laboratories). For analysis of newly synthesized DNA, trypanosomes were collected by centrifugation at 1,000 g for 10 min, re-suspended in thymidine free HMI-11, and induced with tetracycline as required 24 h later. 5-ethynyl-2′-deoxyuridine (EdU) (37) incorporation was assessed as previously described (38), using the Click-iT EdU Alexa Fluor 555 imaging kit (Life Technologies), with some modifications: briefly, cells were incubated with 150 µM EdU for 4-6 h. EdU was washed off and cells were re-suspended in 500 µl of media and mixed 1:1 with 2 % formaldehyde in PBS for 1 h at 4 °C. The formaldehyde was removed by washing cells twice in PBS and re-suspending in 1 % BSA in dH_2_O before spreading on glass slides and drying overnight. Cells were re-hydrated in PBS and the azide click chemistry reaction was performed as per the manufacturer’s instructions. All cells were mounted in Vectashield containing DAPI and images were acquired using an Axiovert 200M fluorescent microscope (Zeiss) coupled to an Axiovert MRm camera. Images were then analysed using Fiji (36).

### Flow cytometry

For cell cycle analysis, cells were collected by centrifugation (1,000 g, 10 min) and washed in ice cold PBS before being re-suspended in 300 µl ice cold PBS and fixed overnight with 700 µl methanol at -20 °C. Fixed cells were washed twice with PBS before DNA staining with propidium iodide at 5 µg.ml^−1^, and RNA digestion with RNase A at 10 µg.ml^−1^ for 1 hour at 37 °C. For VSG detection, the primary antibodies were rat α-VSG-2 (1:10,000) and rabbit α-VSG-5 (1:10,000). Secondary antibodies were goat α-rat Alexa Fluor 647 (1:2,000) and goat α-rabbit Alexa Fluor 488 (1:2,000). Samples were analyzed on a BD FACSCanto (BD Biosciences), and data were visualized and processed using FlowJo software.

### RNA-seq

RNA-seq was performed as described previously (39). Briefly, 2T1, MCP*^VSG-2^* or RNAi*^VSG-2^*cells were incubated with tetracycline for 0, 8 and 12 h. 1 × 10^8^ cells per condition were washed with PBS and total RNA extracted using an RNeasy kit (QIAGEN) according to the manufacturer’s instructions. Samples were sequenced in triplicate on a DNBSEQ-G400 platform (BGI, Hong-Kong). Raw sequencing data was processed through a standardized pipeline. initial quality control was performed using FastQC (https://www.bioinformatics.babraham.ac.uk/projects/fastqc/), followed by adapter trimming and quality filtering with Fastp (0.20.0) (40). Processed reads were aligned to the *T*. *brucei* TREU927 reference genome v68 (41) supplemented with a set of 1,200 bp truncated *VSGs* using Bowtie2 (2.3.5) (42) with ‘--very-sensitive-local’ parameters. The resulting alignments were processed with SAMtools (1.9) (43) for sorting and indexing, and PCR duplicates were marked using Picard MarkDuplicates (2.22.3) (44). Read counts per coding sequence were quantified using featureCounts (1.6.4) (45) with parameters: -p (pair end) -B (both ends successfully aligned) -C (skip fragments that have their two ends aligned to different chromosome) -M (count multi-mapping) -O (match overlapping features) -t CDS (count level) -g gene_id (summarization level). Genes with low counts were filtered out using edgeR (46). Overall quality metrics for fastq files and alignments were aggregated and visualized with MultiQC (47). Differential abundance analyses were carried out in R (3.6.1) with edgeR (3.28.0) using generalized linear models (GLM) and the correction factors for length and GC bias provided by the cqn package (1.32.0) (47).

### Proteomics

Mass spectrometry and proteomic analyses were performed as described previously (16). MCP*^VSG-2^* or RNAi*^VSG-2^* cells were grown for 24 h with or without tetracycline. 5 × 10^7^ cells were washed in PBS and resuspended in 100 μL of a solution containing 5 % SDS and 100 mM triethylammonium bicarbonate. Triplicate samples were submitted to the Fingerprints Proteomics Facility at the University of Dundee for analysis. Cell lysates were treated with 25 U of Benzonase (EMD Millipore, #70664) and sonicated for 2 minutes in a waterbath sonicator. The protein concentration was then determined using the Micro BCA P3rotein Assay Kit (Thermo Fisher, #23235). From each lysate, a volume equivalent to 150 µg of protein was processed using S-Trap mini spin columns (Protifi, # CO2-mini-80) and following the default protocol. Briefly, the lysates were reduced and alkylated by the addition of 20 mM dithiothreitol (VWR, #M109-5G) and 40 mM Ioadoacetamide (Sigma-Aldrich, #I6125-10G), respectively. The proteins were then precipitated with the addition of 12 % Orthophosphoric acid (VWR, #20624.262) and 7x sample volume of Strap Binding Buffer (90 % methanol (VWR, #83638.290) containing 100 mM TEAB (Sigma-Aldrich, #T7408-100ML). The acidified mixture was then placed into the spin columns, and after a centrifugation step, the columns were washed with Strap Binding Buffer. The proteins were digested overnight with the addition of trypsin (1:40, Thermo Fisher, #90057) at 37 °C in a water-saturated atmosphere. Fresh trypsin (1:40) was then added and incubated for a further 6 h. The peptides were then eluted from the columns by adding 50 mM TEAB and centrifuging for 30 sec; two more elution steps using 0.2 % aqueous formic acid (Fisher Chemical, #A117-50), and 50 % aqueous acetonitrile (VWR, #83640.290) containing 0.1 % formic acid were also carried out. The peptides were then dried by vacuum centrifugation. The dried peptides were resuspended with 20 µl of 1 % formic acid and were injected into a Q-Exactive Plus Mass Spectrometer (Thermo Fisher) for quality control assessment and quantification. Samples volumes equivalent to 1.5 µg of peptides were injected onto a nanoscale C18 reverse-phase chromatography system (Ultimate 3000 RSLC nano, Thermo Scientific) and elctrosprayed into an Orbitrap Exploris 480 Mass Spectrometer (Thermo Fisher). For liquid chromatography, the following buffers were used: Buffer A (0.1 % Formic Acid (FA, Fisher Scientific, #A117-50) in MilliQ water (v/v)) and Buffer B (80 % acetonitrile (VWR, #83640.290, 0.1 % FA in MilliQ water (v/v)). Samples were loaded at 10 µl/min onto a trap column (100 µm x 2 cm, PepMap nanoViper C18 column, 5 µm, 100 Å, Thermo Scientific) equilibrated with 0.1 % trifluoroacetic acid (TFA, Thermo Scientific, #85183). The trap column was washed for 3 min at the same flow rate with 0.1 % TFA then switched in-line with a Thermo Scientific, resolving C18 column (75 μm × 50 cm, PepMap RSLC C18 column, 2 μm, 100 Å). Peptides were eluted from the column at a constant flow rate of 300 nl/min with a linear gradient from 3 % buffer B to 6 % buffer B in 5 min, then from 6 % buffer B to 35 % buffer B in 115 min, and finally from 35 % buffer B to 80 % buffer B within 7 min. The column was then washed with 80 % buffer B for 4 min. Two blanks were run between each sample to reduce carry-over. The column was kept at 50 °C. The data was acquired using an easy spray source operated in positive mode with spray voltage at 2.40 kV, and the ion transfer tube temperature at 250 °C. The MS was operated in DIA mode. A scan cycle comprised a full MS scan (m/z range from 350-1650), with RF lens at 40 %, AGC target set to custom, normalised AGC target at 300 %, maximum injection time mode set to custom, maximum injection time at 20 ms, microscan set to 1 and source fragmentation disabled. MS survey scan was followed by MS/MS DIA scan events using the following parameters: multiplex ions set to false, collision energy type set to normalized, HCD collision energies set to 25.5, 27 and 30 %, orbitrap resolution 30000, first mass 200, RF lens 40 %, AGC target set to custom, normalized AGC target 3000 %, microscan set to 1 and maximum injection time 55 ms. Loop control N, N (number of spectra set to 23). Data for both MS scan and MS/MS DIA scan events were acquired in profile mode. Analysis of the DIA data was carried out using Spectronaut (version 17.4.230317.55965, Biognosys, AG). The directDIA workflow, using the default settings (BGS Factory Settings) with the following modifications was used: decoy generation set to inverse, Protein LFQ Method was set to QUANT 2.0 (SN Standard) and Precursor Filtering set to Identified (Qvalue); Precursor Qvalue Cutoff and Protein Qvalue Cutoff (Experimental) set to 0.01; Precursor PEP Cutoff set to 0.01, Protein Qvalue Cutoff (Run) set to 0.01 and Protein PEP Cutoff set to 0.75. Cross-run normalization was selected, with Normalization Strategy set to Global (normalizing on the median). For the Pulsar search the settings were: maximum of 2 missed trypsin cleavages; PSM, Protein and Peptide FDR levels set to 0.01; scanning ranges et to 300-1800 m/z and Relative Intensity (Minimum) set to 5 %; cysteine carbamidomethylation set as fixed modification and acetyle (N-term), damication (asparagine, glutamine), dioxidation (methionine, tryptophan), glutamine to pyro-Glu and oxidation of methionine set as variable modifications. Searches were made using a protein database of *T. brucei* TREU927 v51 obtained from TriTrypDB (41) combined with a predicted set of VSG proteins from the Lister 427 strain truncated at the first 400 AA.

### Differential protein abundance

Data analysis was performed using custom Python and R scripts, using the SciPy ecosystem of open-source software libraries (48). A protein group pivot table was exported from the output of the Spectronaut analysis v15 (Biognosys). The protein groups identified as single hits were considered missing values. Protein groups with missing values in more than 50 % of the samples were excluded from the analysis. The differential expression analysis was performed with limma v3.54 (49) after log2 transformation of the data. FDR values were computed with the toptable function in limma.

## RESULTS

### Blocking *VSG* translation via MCP recruitment to the 5’-UTR

To probe roles for the *VSG* transcript that extend beyond encoding VSG, we sought a method to establish bloodstream-form trypanosomes in which *VSG* translation was specifically and conditionally blocked. For this purpose, we selected a system comprising the bacteriophage MS2 coat protein (MCP), which binds the MS2 RNA hairpin. This approach has been widely used in other organisms to block translation (35) and also in *T. brucei* to block GFP expression (50). Morpholino oligonucleotides can also be used to block translation and have been used to block *VSG* translation (51), but this approach typically lacks conditional regulation and was not considered sufficiently penetrant for the studies we proposed.

Although *T. brucei* cells are diploid, a single *VSG* gene is expressed at a hemizygous sub-telomeric locus (52). To block translation of the single expressed *VSG-2* gene, we first assembled a strain for tetracycline-inducible expression of MCP^GFP^; with an *N*-terminal La nuclear localization signal (33), and with GFP fused to the C-terminus. MCP^GFP^ expression was tetracycline-inducible (Fig. 1A), and the protein accumulated in the nucleus, as expected (Fig. 1B). We next generated strains containing both inducible MCP^GFP^ and a single MS2 ‘hairpin-sequence’ in the 5’-UTR of the *VSG-2* gene (Fig. 1C). Correct integration of the hairpin-sequence in the *VSG-2* gene, and incorporation into *trans*-spliced mRNA in these ‘MCP*^VSG-2^*’ cells was confirmed by RT-PCR and sequencing (Fig. 1C).

**Figure 1.**
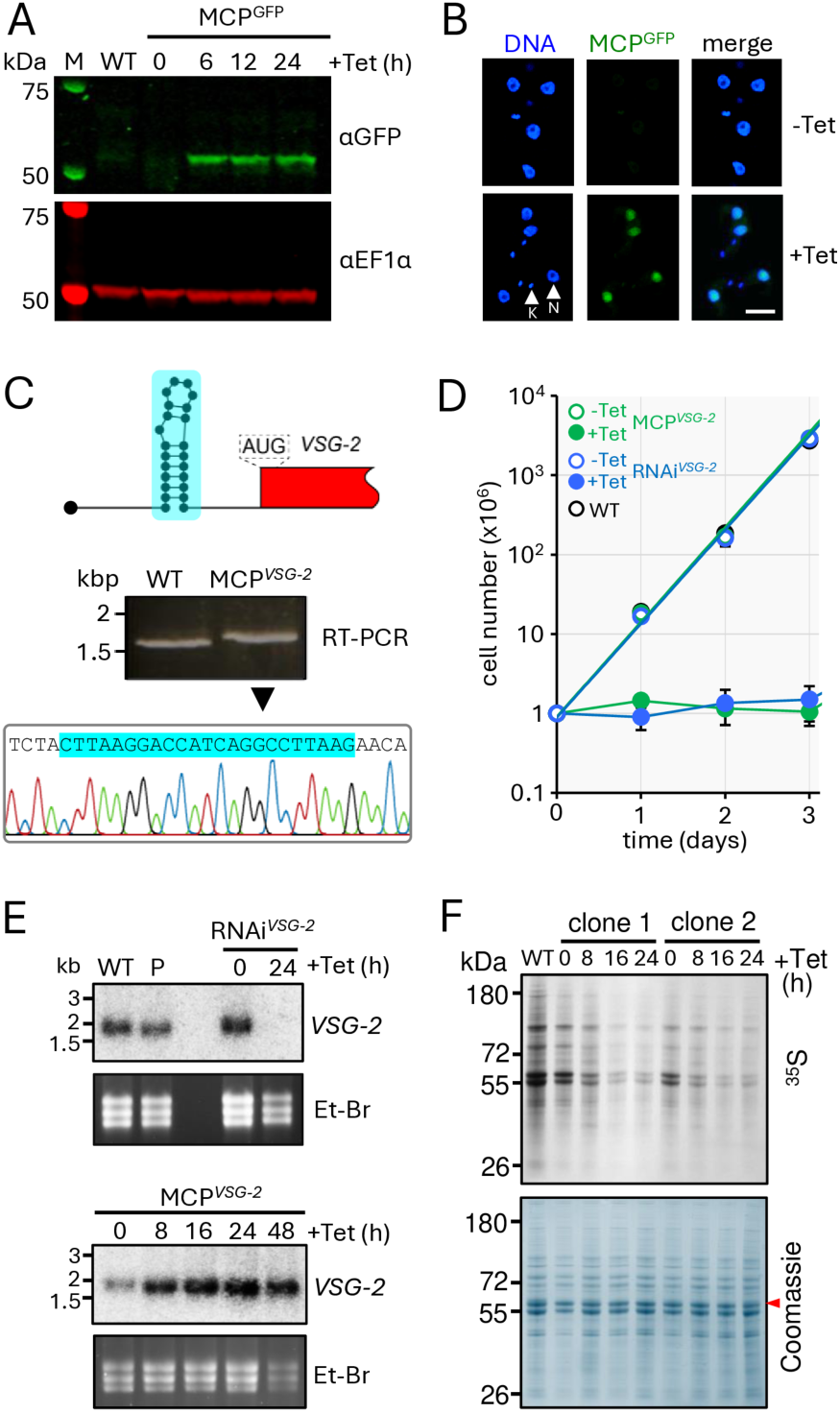
Blocking *VSG* translation via MCP recruitment to the 5’-UTR. (**A**) The protein blot shows tetracycline-inducible expression of MCP^GFP^ in *T. brucei*. EF1-α serves as a loading control on the same blot. (**B**) Fluorescence microscopy reveals MCP^GFP^ in *T. brucei* nuclei following induction (+Tet, 24 h). Nuclear (N) and mitochondrial kinetoplast (K) DNA are indicated. (**C**) The schematic indicates an MCP-binding, MS2-RNA hairpin sequence in the *VSG-2* 5’-UTR in MCP*^VSG-2^* cells. The gel shows products obtained after *VSG-2*-specific RT-PCR, and the sequence trace shows incorporation of the MS2 hairpin sequence in MCP*^VSG-2^* cells. WT, wild-type. (**D**) Cumulative growth curves for wild-type cells and MCP*^VS-2^* and RNAi*^VSG-2^* cells before and after induction. The data represent averages from two independent biological replicates. (**E**) The RNA blots show *VSG-2* mRNA abundance following knockdown in RNAi*^VSG-2^* cells or following MCP^GFP^ induction in MCP*^VSG-2^* cells. Ethidium bromide (Et-Br) stained gels serve as loading-controls. The data are representative of two independent biological replicates. WT, wild-type, P, parental strain. (**F**) Metabolic labeling with ^35^S methionine during MCP^GFP^ induction in MCP*^VSG-2^* cells; both biological replicates. The Coomassie stained panel serves as a loading control; the red arrowhead indicates VSG-2.

For comparison with MCP-based translation blockade, we generated RNAi*^VSG-2^* strains containing an inducible *VSG-2* knockdown cassette. We then compared growth of wild-type cells, and the MCP*^VSG-2^*and RNAi*^VSG-2^* strains under non-inducing and inducing conditions. All strains displayed comparable exponential growth under non-inducing conditions (Fig. 1D), indicating both that MCP^GFP^ expression is tightly regulated and that the modified *VSG-2* mRNA had no detectable deleterious impact on fitness in the MCP*^VSG-2^* strains prior to MCP^GFP^ induction. Following induction of knockdown, the RNAi*^VSG-2^* strains displayed a severe growth defect under inducing conditions, as expected (53), and the MCP*^VSG-2^*strains displayed a similar profile following induction of MCP^GFP^ expression (Fig. 1D).

Using RNA blotting, we next assessed *VSG-2* transcript abundance following induction of *VSG-2* knockdown or translation blockade. This analysis confirmed *VSG-2* mRNA knockdown in RNAi*^VSG-2^* cells as anticipated (Fig. 1E). In contrast, *VSG-2* mRNA was stabilized following induction of MCP^GFP^ expression in MCP*^VSG-2^* cells; indeed *VSG-2* mRNA abundance was increased in these cells (Fig. 1E).

To determine whether *VSG* translation was indeed blocked in MCP*^VSG-2^* cells, we analyzed protein synthesis under inducing conditions. Metabolic labeling of newly synthesized proteins with ^35^S-methionine revealed reduced global translation in two independent MCP*^VSG-2^* strains (Fig. 1F) and quantification revealed that protein synthesis was reduced by 90 % after 16 h. Thus, VSG translation blockade brings about global translation arrest, as also reported following *VSG* RNAi (54). We conclude that MCP^GFP^-dependent *VSG* translation blockade, like *VSG* transcript knockdown, triggers global translation arrest and a severe growth defect. VSG translation blockade therefore phenocopies defects associated with loss of VSG expression, while maintaining the cellular pool of *VSG* mRNA. The major differences in *VSG-2* mRNA abundance observed in MCP*^VSG-2^* and RNAi*^VSG-2^* strains, therefore, presented an opportunity to investigate specific roles of the *VSG* mRNA.

### Native *VSG* exclusion is sustained following VSG-2 perturbations

To investigate specific non-coding roles for the *VSG* transcript, we first used RNA-seq to assess the transcriptomes following either *VSG-2* knockdown or translation blockade. We induced each perturbation for either 8 h or 12 h. This analysis confirmed *VSG-2* knockdown in RNAi*^VSG-2^* cells and increased *VSG-2* transcript abundance following translation blockade in MCP*^VSG-2^* cells (Fig. 2), supporting the results from RNA blotting above (Fig. 1E). Knockdown reduced *VSG-2* transcript by >90 % while translation blockade increased the abundance by >2-fold, confirming that translation blockade following recruitment of the MCP does indeed stabilize the *VSG-2* transcript. Although VSG feedback to regulate *VSG* transcript abundance has been reported previously (55), this increase in abundance of the already super-abundant *VSG-2* transcript is quite remarkable, given a starting-point of approximately 10 % of total cell mRNA (9). We conclude that *VSG-2* transcript abundance was 24 or 39-fold lower in RNAi*^VSG-2^* cells than in MCP*^VSG-2^* cells following 8 or 12 h of induction, respectively.

**Figure 2.**
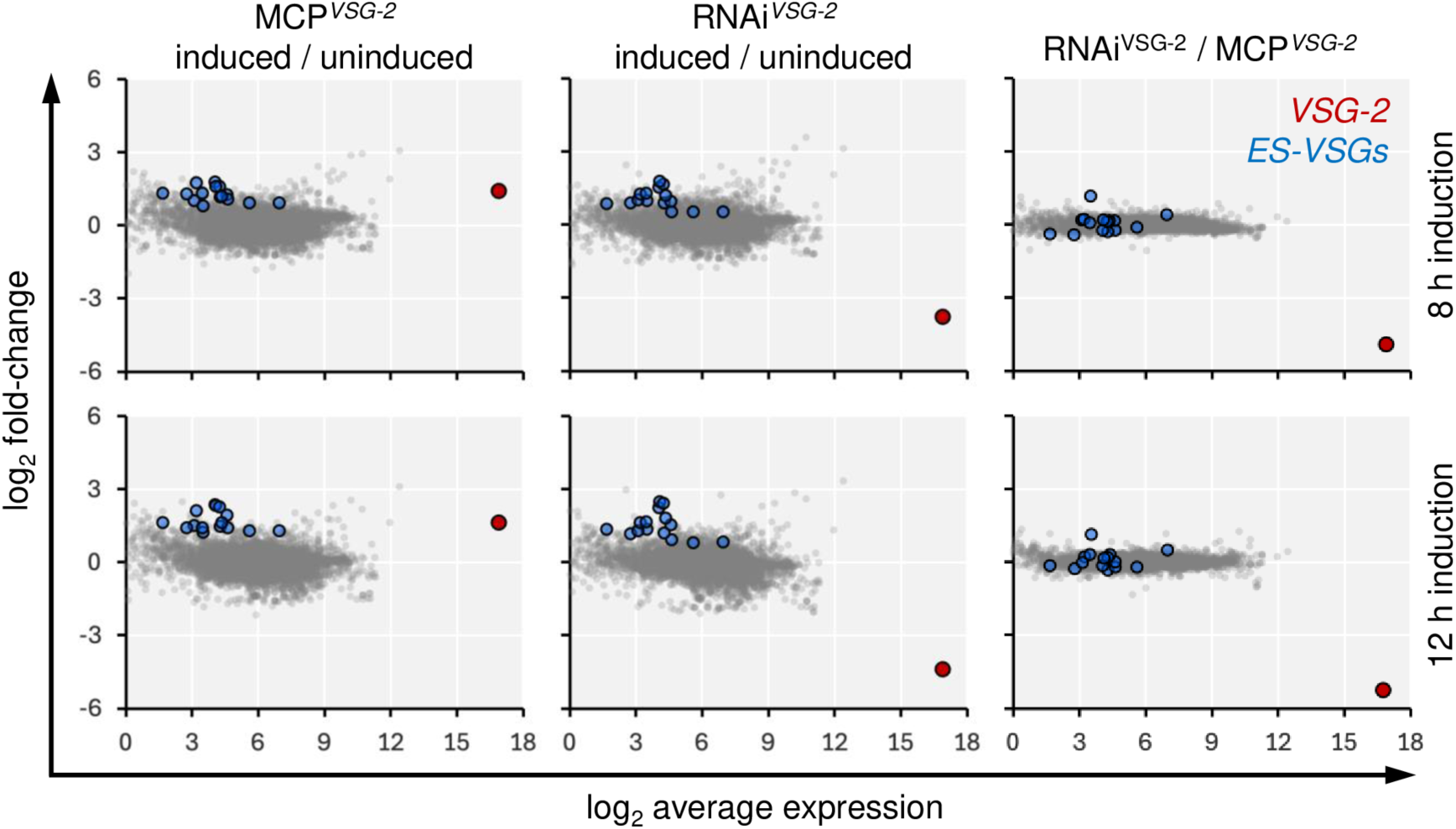
Native *VSG* exclusion is sustained following VSG-2 perturbation. RNA-seq analysis following induction of *VSG-2* mRNA knockdown in RNAi*^VSG-2^* cells or translation blockade in MCP*^VSG-2^* cells for either 8 h (upper panels) or 12 h (lower panels). *VSG-2* and other excluded expression-site associated *VSG* transcripts are highlighted. The data represent averages from three independent technical replicates in each case. n = 8458.

Despite the difference in *VSG-2* transcript abundance, other telomeric *VSGs* were not substantially de-repressed in either of these strains (Fig. 2). Among fifteen expression-site associated *VSGs* detected, we saw <3-fold average de-repression in RNAi*^VSG-2^* and MCP*^VSG-2^*strains, with these *VSGs* remaining >2,500-fold lower in abundance on average than the unperturbed active *VSG-2* transcript. Indeed, with the notable exception of *VSG-2,* the transcriptomes of RNAi*^VSG-2^*and MCP*^VSG-2^* strains were otherwise similar following 12 h of induction (Fig. 2, right-hand panels). Thus, neither knockdown of the active *VSG* transcript using RNA interference, nor blocking translation by recruiting MCP, substantially impacted the expression of established excluded *VSGs*. We conclude that *VSG* exclusion was sustained when the dominant *VSG* transcript was depleted for 12 h, and by >15-fold.

### A *VSG* transgene evades exclusion when the *VSG-2* transcript is depleted

The results above indicated that native *VSG* exclusion was maintained when *VSG-2* mRNA was depleted or when *VSG-2* translation was blocked. Next, we asked whether *VSG* transcripts compete for establishment of the dominant active state. To address this question, we used an expression assay with a *VSG* transgene that is known to be subject to exclusion (19). MCP*^VSG-2^*and RNAi*^VSG-2^* strains were transiently induced for three hours prior to washing and delivery of the *VSG-5* transgene which, when integrated into the genome, is transcribed from an *rDNA* promoter adjacent to a *de novo* telomere (19). Five days following delivery of the transgene, the resulting cells were stained for cell-surface VSG-2 and VSG-5 expression and were assessed using immunofluorescence microscopy and flow cytometry (Fig. 3A).

**Figure 3.**
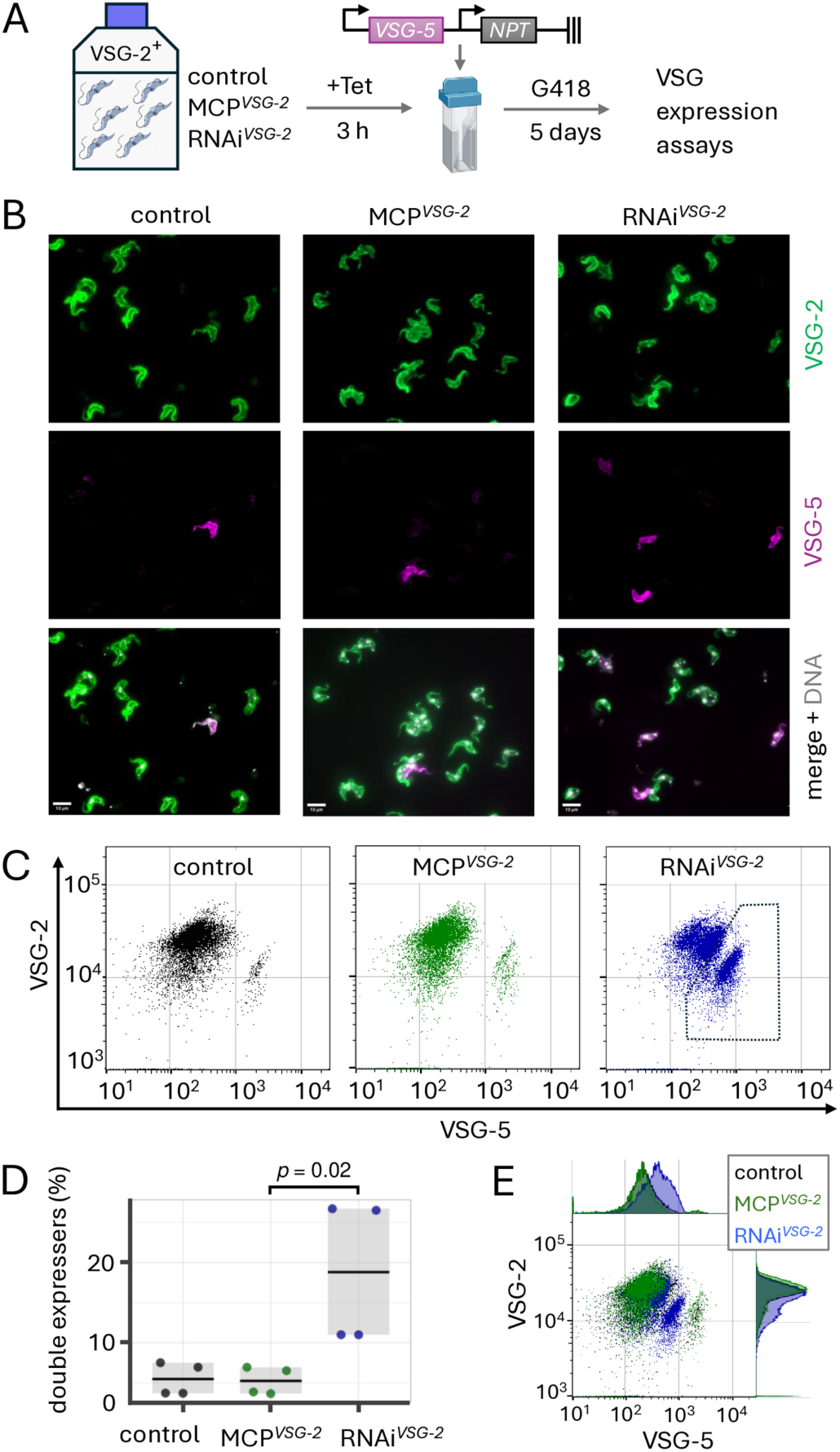
A *VSG* transgene evades exclusion when *VSG-2* is depleted. (**A**) the schematic illustrates the experimental procedure. MCP*^VSG-2^*and RNAi*^VSG-2^* strains were induced with tetracycline (Tet) for 3 h, Tet was removed, and cells were transfected with the *VSG-5* reporter; the arrows indicate *rDNA* promoters, the symbol on the right indicates a *de novo* telomere. Cells were then selected for *NPT* expression using G418 and *VSG* expression was assessed after 5 days. (**B**) Representative fluorescence microscopy images reveal VSG-2 and VSG-5 expression. Scale bars, 10 μm. (**C**) Flow cytometry analysis of VSG-2 and VSG-5 expression. n = 10,000 cells in each case. (**D**) Quantification of cells expressing both VSG-2 and VSG-5 according to the gated region indicated in C (right-hand panel). The data represent averages from two independent biological replicates and two technical replicates. Horizontal lines indicate mean values. (**E**) Data from C were combined to compare bulk population profiles for each strain.

The *VSG-5* transgene behaved as expected in the control cells (19), with only 5.5 % of cells on average expressing the transgene, and the results were very similar (5.2 %) following transient *VSG*-2 translation blockade in MCP*^VSG-2^* cells (Fig. 3B-C, left-hand and middle panels; Fig. 3D-E). In contrast, the VSG-5 signal was increased for a significantly higher proportion of cells (18.8 %) following transient *VSG*-2 knockdown (Fig. 3B-C, right-hand panels; Fig. 3D-E). Indeed, the bulk population of RNAi*^VSG-2^* cells displayed increased VSG-5 expression relative to control and MCP*^VSG-2^*cells (Fig. 3E); similar results were obtained in three separate experiments conducted by two independent investigators. We conclude that a transgenic *VSG-5* reporter evaded exclusion when delivered under transient *VSG-2* knockdown. These results suggest that the establishment of monogenic *VSG* expression is driven by competing *VSG* transcripts.

### *VSG* transcript perturbation impacts CFB2 abundance

We next wondered whether phenotypes associated with *VSG* perturbation might be associated with specific changes in VSG or regulatory factor abundance, and we used quantitative proteomic analysis to address this question. Consistent with failure to synthesize new VSG, we observed similar and significant (>30 %, *p* = 4.2e-7) VSG-2 depletion following 24 h of either *VSG-2* translation blockade or knockdown (Fig. 4A); depletion was likely limited because VSG has a half-life of approx. 30 h (56). As observed using transcriptome analysis above (Fig. 2), other expression-site associated *VSGs* were moderately de-repressed in both the MCP*^VSG-2^* and RNAi*^VSG-2^* strains (Fig. 4A); although those VSGs detected remained >500-fold lower in abundance on average than unperturbed VSG-2. Notably though, we did observe significantly higher, albeit <2-fold on average, VSG de-repression following knockdown (Fig. 4B).

**Figure 4.**
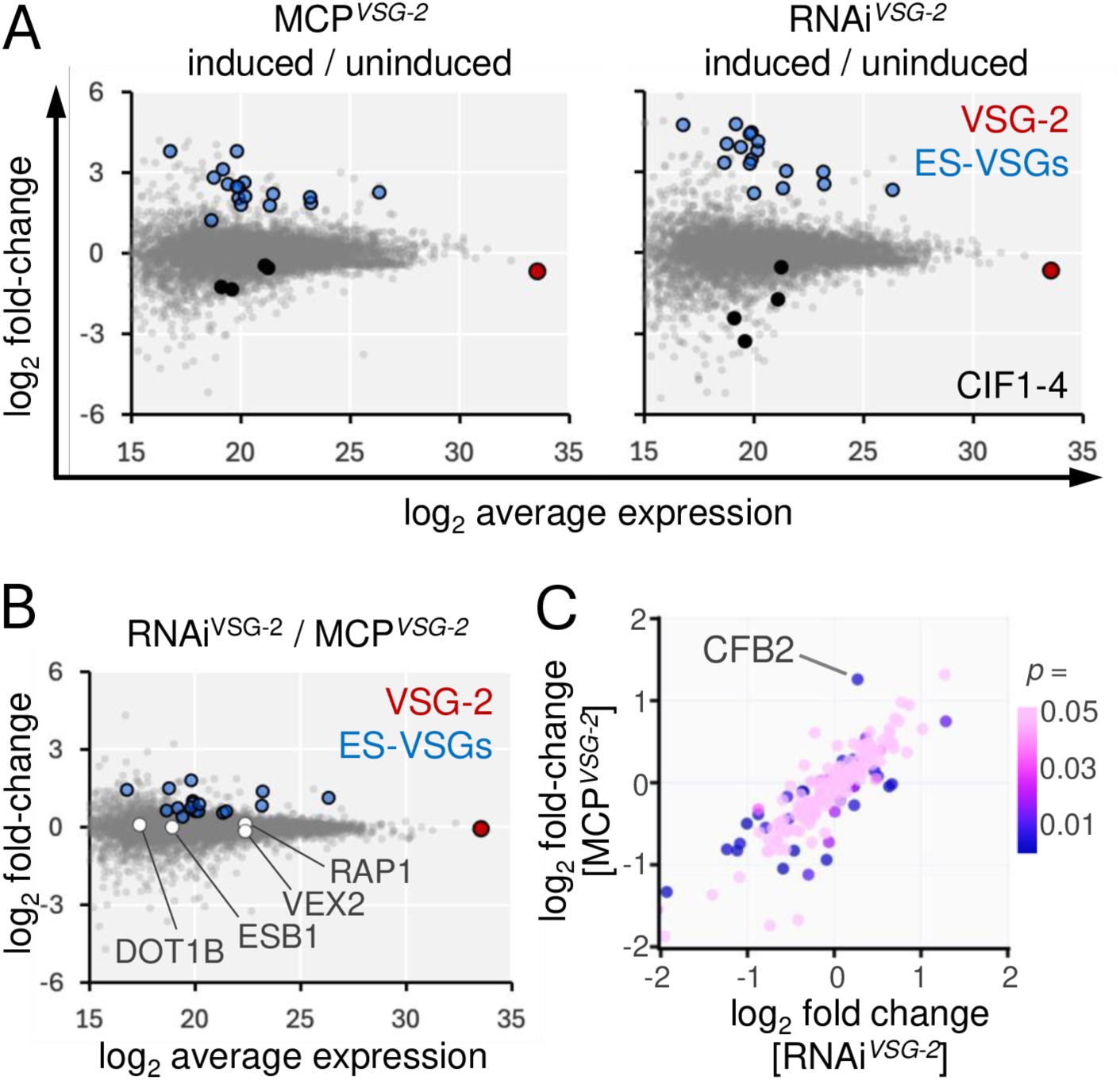
*VSG* transcript perturbation impacts CFB2 abundance. (**A**) Proteomics analysis following induction of *VSG-2* mRNA knockdown in RNAi*^VSG-2^* cells or translation blockade in MCP*^VSG-2^* cells for 24 h. *VSG-2*, other excluded expression-site associated *VSG* transcripts, and CIF1-4 are highlighted. The data represent averages from three independent technical replicates in each case. n = 5552. (**B**) As in A, but showing a comparison between *VSG-2* mRNA knockdown in RNAi*^VSG-2^* cells and translation blockade in MCP*^VSG-2^*cells. *VSG* regulatory factors are also highlighted. (**C**) Proteomics analysis of all detected proteins associated with the ‘mRNA-binding’ Gene Ontology term (n = 181). The cyclin-like F-box protein, CFB2 was significantly increased in abundance (*p* = 0.004) following *VSG-2* translation blockade in MCP*^VSG-2^* cells.

The proteomes of MCP*^VSG-2^* and RNAi*^VSG-2^*strains were otherwise similarly perturbed following induction (Fig. 4B). For example, Gene Ontology analysis of the 200 proteins most significantly reduced in abundance in each case revealed enrichment for ‘ribosome biogenesis’ (RNAi*^VSG-2^*, *p* = 1e-5; MCP*^VSG-2^ p* = 3.7e-17) and ‘cleavage furrow’ (RNAi*^VSG-2^*, *p* = 1.4e-4; MCP*^VSG-2^ p* = 2.8e-3), consistent, respectively, with the translation blockade and cytokinesis arrest phenotypes described above. Indeed, all four components of the cytokinesis initiation factor (CIF1-4) (57) were significantly reduced in abundance in both strains (Fig. 4A). We next assessed several known *VSG* expression regulators for differences in abundance following either perturbation. This analysis revealed no significant differences for ESB1, VEX2, RAP1 or DOT1B (Fig. 4B), but did reveal a significant difference for CFB2 (*p* = 4.4e-3). Indeed, analysis of all proteins associated with the ‘mRNA-binding’ Gene Ontology term (n = 181) highlighted CFB2 (Fig. 4C), which was increased in abundance (>2-fold) following translation blockade, consistent with binding and stabilization of the *VSG* transcript (17), but was not significantly changed following *VSG-2* knockdown.

### An additional round of DNA replication in the presence of the *VSG* transcript

Although it is known that *VSG* knockdown leads to S phase, mitosis and cytokinesis arrest (53,58), potential roles for the *VSG* transcript in controlling progression through the cell cycle have not been explored. Our MCP*^VSG-2^* and RNAi*^VSG-2^*strains presented an opportunity to explore such roles. Indeed, we were particularly interested in exploring connections between *VSG* transcript and DNA replication control since the transcript binds CFB2, which also interacts with the S-phase kinase associate protein, SKP1 (17). To assess the impact of translation blockade or *VSG-2* knockdown on cell cycle progression, we used DNA staining and microscopy to visualize nuclear and mitochondrial (kinetoplast) DNA. This analysis revealed a similar dramatic increase in post-mitotic (2N:2K) cells after only 8 h of induction in both cases (Fig. 5A). A striking difference, however, was the emergence of cells with supernumerary (>2) nuclei, specifically following translation blockade, which comprised >40 % of the population 24 h after induction, and >80 % at 48 h (Fig. 5A). These results are consistent with the view that efficient VSG trafficking to the cell-surface is required for cytokinesis (53), and also now indicate that retention of the *VSG* transcript allows an additional round of mitosis.

**Figure 5.**
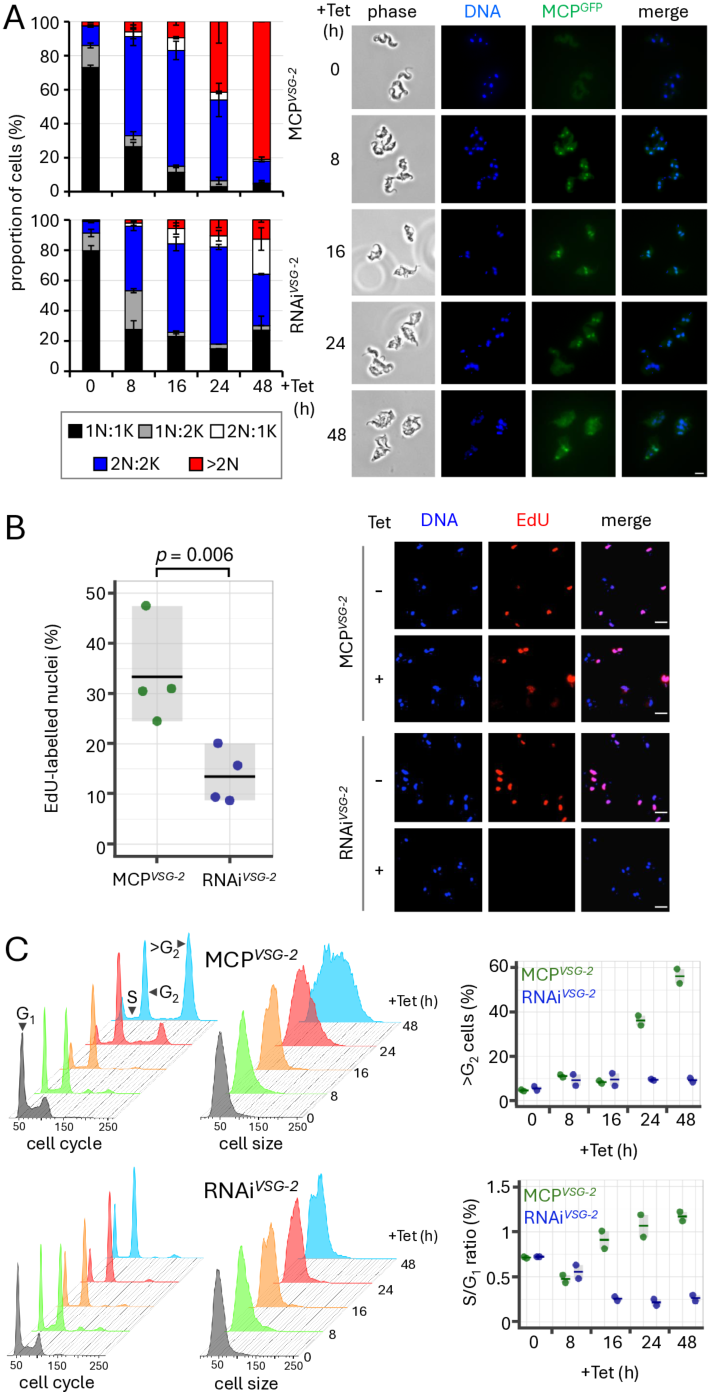
An additional round of DNA replication in the presence of the *VSG* transcript. (**A**) Microscopy-based quantification of the number of nuclei and kinetoplasts per cell following induction in MCP*^VSG-2^* cells (upper panel) or RNAi*^VSG-2^* cells (lower panel). The data represent averages from two independent biological replicates. Representative fluorescence microscopy images of MCP*^VSG-2^* cells are shown on the right. 2N cells (white and blue categories) are post-mitotic. DNA was stained with DAPI while GFP reveals the expression and location of MCP^GFP^. Scale bar, 5 µm. (**B**) Microscopy-based quantification of the proportion of EdU-labelled nuclei in MCP*^VSG-2^* or RNAi*^VSG-2^* cells following induction for 16 h. The data represent averages from two independent biological replicates assessed in duplicate. Horizontal lines indicate mean values. Representative fluorescence microscopy images are shown on the right. DNA was stained with DAPI. Scale bars, 10 µm. (**C**) Flow cytometry analysis following MCP^GFP^ induction in MCP*^VSG-2^* cells, or *VSG-2* knockdown in RNAi*^VSG-2^* cells. Representative histograms indicate DNA content based on propidium iodide-staining (left-hand “cell cycle” panels) or relative cell size based on side-scatter (right-hand “cell size” panels). The plots on the right indicate proportions of cells with a >G_2_ DNA content and the S phase to G_1_ ratios. The data represent averages from two independent biological replicates. Horizontal lines indicate mean values.

To explore DNA replication status, we labelled cells for 6 h with 5-ethynyl-2’-deoxyuridine (EdU) and quantified the proportion of cells replicating their nuclear genome by fluorescence microscopy. DNA in control cells, and in uninduced MCP*^VSG-2^* and RNAi*^VSG-2^* cells (Fig. 5B), was efficiently labeled with EdU (94, 98 and 97 % of cells, respectively). Although both MCP*^VSG-2^* and RNAi*^VSG-2^* strains displayed decreased EdU-labeling following growth under inducing conditions, labeling was significantly higher (*p* = 0.006) in the MCP*^VSG-2^* population, than in the RNAi*^VSG-2^*population (Fig. 5B). Finally, we used flow cytometry to quantify DNA content. Consistent with the DNA-staining, EdU-labeling, and microscopy analysis (Fig. 5A-B), we observed a specific increase in cells with a >G_2_-phase DNA content, and an increase in cell size, in the induced MCP*^VSG-2^*cells (Fig. 5C), indicating endoreduplication in cells that retain the *VSG* transcript. Although pre-cytokinesis arrest progressively diminishes the proportion of G_1_ cells in both cases, the S phase/G_1_ ratio was significantly different 16 h post-induction (*p* = 0.02) and thereafter, being increased in the induced MCP*^VSG-2^* cells and decreased in induced RNAi*^VSG-2^* cells (Fig. 5C). Thus, we observed an additional round of DNA replication and mitosis in the presence of the *VSG* transcript, that was not observed following loss of the *VSG* transcript.

## DISCUSSION

To probe roles of the *VSG* transcript that extend beyond encoding VSG, we assembled and compared strains in which *VSG* translation was conditionally blocked, with strains in which *VSG* transcript was conditionally knocked down. The *VSG* transcript was found to be required to establish silencing of a *VSG* transgene and was also linked to DNA replication control. We suggest that *VSG* transcripts drive competition among *VSG* alleles to establish dominance and exclusion.

Blocking VSG synthesis triggers a pre-cytokinesis arrest (51,53), and a global translation arrest (54) in bloodstream form *T. brucei*, and similar phenotypes are observed following knockdown of other factors with roles in VSG coat maintenance; PFR2 (paraflagellar rod 2), actin and clathrin (59–61), for example. We now confirm that the presence of the *VSG* transcript fails to rescue these phenotypes. On the other hand, our findings suggest that the *VSG* transcript can promote DNA replication, perhaps via a mechanism involving binding the bloodstream form specific cyclin-like F-box protein, CFB2 (16,17). Metazoan Y RNAs have also been implicated in promoting DNA replication (62,63), while embryonic stem cells can proliferate independent of G_1_ cyclins (64). In addition, mammalian long noncoding RNAs regulate the expression of cyclins and CDKs (65); the *gadd6* lncRNA regulates the G_1_/S checkpoint by promoting the degradation of the *Cdk6* transcript (66), for example.

There are remarkable differences in cell cycle controls operating in bloodstream and insect-form *T. brucei*. CRK1 and CRK2 promote the G_1_/S transition in insect-form cells but are not required for this purpose in bloodstream-form cells (67), for example. DNA replication continues in bloodstream-form cells, but not in insect-form cells, following knockdown of the CRK3 partner, CYC6 (68), or the chromosomal passenger protein, aurora-B kinase, AUK1 (69). Knockdown of flagellar function (70), actin (61) or glycophosphatidylinositol (GPI) anchor biosynthesis (71), all of which perturb VSG coat maintenance in bloodstream-form *T. brucei*, also result in the accumulation of multi-nucleated cells specifically in the bloodstream-form. Our findings, now connecting the *VSG* transcript to an additional round of DNA replication and mitosis, may explain these bloodstream-form specific endoreduplication phenotypes, and the absence of such a phenotype following *VSG* knockdown (53,58). We suggest that novel checkpoint controls operate in bloodstream-form trypanosomes whereby cells monitor whether the demand for *VSG* mRNA and protein have been satisfied prior to committing to S phase or cytokinesis, respectively.

We propose a form of RNA-mediated silencing in the *VSG* exclusion system. Indeed, we previously suggested a role for RNA-based, homology-dependent repression in *VSG* allelic exclusion, since reporters lacking *VSG*-associated sequences were also subject to exclusion (19). RNA interference operates in *T. brucei*, but argonaut 1 (AGO1) knockout had no impact on *VSG* exclusion (72), suggesting an RNAi-independent mechanism. Regulation of other genes by anti-sense RNAs that modulate translation (73), or a long non-coding RNA that promotes differentiation (74), have also been reported in *T. brucei*. Specific sequences in the *VSG* transcript may be involved in the non-coding functions we propose here. In this regard, it is notable that the 3’-UTR contains a highly conserved 16-mer motif immediately preceding the polyadenylation site. This motif is thought to bind CFB2 (17), RAP1 (20), and also to be required for m^6^A modification (27). The transgene we used in our assays incorporated a 76 b 3′-UTR that is identical to the native active *VSG-2* sequence (19). One possible model involves competition for a limiting *trans*-activator, accompanied by negative control by *VSG* RNA at competing *VSG* loci, via R-loop formation (20,75), for example. Recent findings reported by others are consistent with this model; specifically, transcription of a second *VSG* with a mutated 16-mer failed to silence the active *VSG* (76).

To interpret our findings, it is important to consider distinct mechanisms of establishment and maintenance in the *VSG* exclusion system. We suggest that establishment of exclusion involves a competition among *VSG* transcripts for binding and sequestration of chromatin-associated RNA-binding proteins, such as VEX2 (18), RAP1 (20), or ESB1 (14). Both establishment and maintenance of the active and silent states likely also require the action of additional chromatin-associated factors, including those factors enriched at the active transcription and splicing compartment (12–14). In addition, the DOT1B histone methyltransferase is required to establish the excluded state (23,24), while maintenance of exclusion is compromised when histones are depleted (25). The chromatin chaperone, CAF-1 maintains the VEX-complex at the active site (18,25), while cohesin promotes inheritance of the active *VSG* during S phase (26). Modification of the *VSG* transcript with m^6^A in the polyA-tail (27) may also promote chromatin accessibility (77).

Our findings indicate that the *VSG* transcript is a coding and non-coding RNA (cncRNA), analogous to bi-functional cncRNAs that control developmental processes in vertebrates and plants (78). We propose a “winner-takes-all” model whereby the *VSG* cncRNA competes for sequestration of transactivators, such as ESB1 (14), and VSG exclusion factors (13), thereby modifying nuclear architecture to increase its own transcription and establish dominance. S phase, which may also be subject to control by the *VSG* cncRNA, may be a key point in the cell cycle when competition among *VSG* transcripts operates. Notably, similar competition among transcripts has been proposed to drive olfactory receptor allelic exclusion in mammals (79,80), while ncRNA also impacts *var* gene exclusion in malaria parasites (81). We conclude that the *VSG* transcript is a cncRNA that inhibits the expression of its competitors.

## DATA AVAILABILITY

Sequencing data have been deposited in the European Nucleotide Archive, www.ebi.ac.uk/ena (BioProject ID: PRJNA1216521). The mass spectrometry proteomics data have been deposited at the PRIDE repository (Dataset identifier PXD060524).

## SUPPLEMENTARY DATA

Supplementary Data are available.

## ACKNOWLEDGMENTS

We thank R. Clark for assistance with flow cytometry, A. Score for assistance with proteomics, and N. Zhang and M. Field for advice on metabolic labeling. This work benefitted from the resources provided by VEuPathDB.

## AUTHOR CONTRIBUTIONS

Conceptualization; DOE, SH, JEW, CAM, JRCF, AT, DH. Investigation; DOE, SH, JEW, CAM, JRCF, AT. Data curation and analysis; MT. Supervision; DH. Original draft; DOE, DH. Review & editing; all authors.

